# Improved Source Localization of Auditory Evoked Fields using Reciprocal BEM-FMM

**DOI:** 10.1101/2025.05.09.653081

**Authors:** Derek A. Drumm, Guillermo Nuñez Ponasso, Alexander Linke, Gregory M. Noetscher, Burkhard Maess, Thomas R. Knösche, Jens Haueisen, Jeffrey David Lewine, Christopher C. Abbott, Sergey N. Makaroff, Zhi-De Deng

**Author notes:** First two authors share primary authorship; last two authors share senior authorship (Derek A. Drumm, Guillermo Nuñez Ponasso, Sergey N. Makaroff, Zhi-De Deng).

## Abstract

The precise localization of auditory evoked fields (AEFs) from magnetoencephalography (MEG) data is very important for the functional understanding of the auditory cortex in medicine and cognitive neuroscience. The numerical solution of the field equations in the human head using the boundary element method (BEM) is a powerful tool for achieving this. We hypothesized that the spatial resolution of the BEM is crucial for the achievable accuracy. However, in classical BEM (as implemented, e.g., in MNE-Python), very high resolutions are impractical due to the associated prohibitive computational effort. In contrast, our recently introduced reciprocal boundary element fast multipole method (reciprocal BEM-FMM) allows for hitherto unprecedented spatial resolution. In this work, we apply our reciprocal BEM-FMM technique for source estimation to localize AEFs, and we compare our results with the source estimates produced using a 3-layer BEM model (standard BEM) via MNE-Python. We first validate our methodology through comparison of source estimates of simulated N1m components of AEFs using a receiver operating characteristic (ROC) measure. While we obtain ROC measures of about 80% for the standard BEM, reciprocal BEM-FMM reaches about 90%, a significant statistical improvement. We then apply this methodology to analyze the source estimates of experimental data obtained from a cohort of 7 participants subjected to binaural auditory stimulation. Using a dispersion measurement to quantify the focality of localized sources, we find improvements upwards of 30% using reciprocal BEM-FMM over the standard BEM.

Analyses from both simulated and experimental data show localization of AEFs using high-resolution reciprocal BEM-FMM is significantly better in terms of accuracy and focality than those estimates of the low-resolution standard BEM. We therefore recommend using the high-resolution reciprocal BEM-FMM to utilize high spatial anatomical precision for the modeling of neural activity.

## 1 Introduction

The objective of source localization [19] is to estimate the location of neural activation from non-invasive recordings such as *electroencephalography* (EEG) or *magnetoencephalography* (MEG) [15]. In [28] we introduced a novel forward solver for MEG source localization based on Lorentz’s reciprocity theorem and the *boundary-element fast multipole method* (BEM-FMM) [23, 22]. This new method is based on the computation of cortical basis functions by means of a fictitious current injected through each MEG sensor coil, similar to forward solutions in transcranial magnetic stimulation (TMS). The use of the fast multipole method [13] coupled with the charge-based formulation of the BEM [27] allows us to avoid computing the dense system matrices of the traditional surface-potential BEM [9, 8], and as a result our method enables source localization using multiple high resolution meshes. In [28] we localized somatosensory evoked fields and obtained good results compared to the classical MNE-Python with 3-layer models [11, 12]. In this work, we apply our methods to auditory evoked fields, which are more challenging from a source reconstruction perspective than the somatosensory evoked fields: on the one hand, the binaural auditory stimulation elicits activation on both hemispheres, with two approximately symmetric sources at the far-apart left and right temporal lobes; on the other hand, the head anatomy near the temporal lobes is more complicated than near the parietal lobes, and it is harder to represent accurately using downsampled 3-layer head models. As a result, high-resolution models will be able to provide more accurate source localizations—this was already suggested in the error maps of [29].

Auditory evoked fields (AEFs) are magnetic signals elicited by auditory stimuli. They are localized in the auditory cortex, having components in the Heschl gyri (primary auditory cortex) and in the planum temporale as a secondary area [33, 20, 41, 10]. There are three main components of auditory evoked fields: 1) Middle-latency components, e.g. Pam (¡ 50 ms) which originates in regions medial and lateral to the Heschl gyri; 2) N1m (~100 ms), consistently localized in the lateral Heschel gyri and planum temporale; and 3) P2m (~200 ms), which has been found to have origins near the N1m source locations although with much greater interindividual variability [21].

The source localization of AEFs is relevant from an applications standpoint as they can be used clinically for presurgical mapping in the temporal lobe epilepsy [10], in the identification of functional reorganization in patients with lesions [24], and as a biomarker for language and developmental disorders [31, 32]. The sources of AEF are also very important due to their correlations with mental health and psychological disorders [18, 6, 4].

In this study, we analyze the source localization of auditory evoked fields elicited by binaural auditory stimuli for a cohort of 7 patients with drug-resistant depression, whose data were provided by the authors of the study of [4]. Using each patient’s BEM models, we simulate noiseless N1m peaks of binaural auditory stimulation. We compute the source localization results for both the simulated and experimental N1m peaks. Additionally, we compare the source localization results from our own solver to that of MNE-Python.

## 2 Materials and methods

### 2.1 Forward modeling using reciprocal BEM-FMM

We construct forward models using the reciprocal BEM-FMM approach [28], which hinges on the Lorentz reciprocity theorem relating the primary electric field **E** [V m^−1^] induced by an induction coil outside the cortical surface and the magnetic flux density **B** [T] induced by a current dipole source at location **r** [m] within the cortical surface [34, 17, 26, 28]:

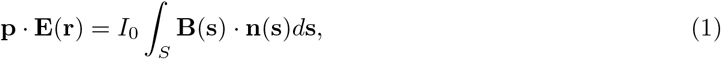

where **p** [A m] is the dipole moment of the current dipole source located at **r**, and *I*_0_ [A] is the amplitude of the current flowing through the induction coil *S*. Assuming the dipole moment is oriented normal to the cortical surface, we can extend Equation 1 to *i* = 1, *…, M* induction coils and *j* = 1, *…, N* current dipoles, yielding

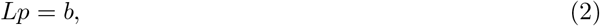

where *L* denotes the lead-field matrix whose entries are given by

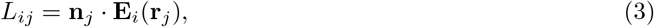

where **E**_*i*_ is the induced electric field of the *i*-th induction coil, *p*_*j*_ is the strength of the *j*-th current dipole, and *b*_*i*_ is the total magnetic flux through the *i*-th induction coil. Equation 2 represents the reciprocal BEM-FMM forward model, with each of the *i* coils acting analogously to TMS coils. However, in the context of MEG source analysis, each of the *i* coils can now be considered as MEG sensors, with the entries of *b* indicating the total measured output of each sensor. In this sense, Equation 2 represents a typical lead-field model of source reconstruction, with *p* denoting the unknown strengths of each of the current source dipoles [19].

Table 1 shows the parameter values used in the BEM-FMM computation of the forward model. Details regarding the forward model computation are found in [28].

**Table 1:** Parameter values used in the computation of the forward model using BEM-FMM. The symbol *µ*_*A*_ denotes the average triangle area in the mesh.

| Parameter | Value |
| --- | --- |
| FMM precision | $1 \times 10^{-4}$ |
| Region of interest radius | $7 \times 10^{-2}[\text{m}]$ |
| Neighbor integral correction radius | $2\sqrt{\mu_A} [\text{m}]$ |
| Characteristic of the coil pulse waveform: $\partial i_j / \partial t$ | $1[\text{A s}^{-1}]$ |

### 2.2 Inverse solutions using minimum norm estimation and dSPM

We compute the maximum a posteriori solution to the inverse problem of solving for source strengths *p* given noisy MEG data *b* [19]. Such solutions are found by using minimum norm estimation to invert Equation 3. This is done using the inverse operator [14]

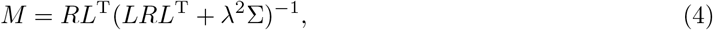

where *R* is the prior source covariance, Σ is the noise covariance, and *λ* is a regularization parameter chosen as the inverse of the signal-to-noise ration, *λ* = 1/SNR. In this way, the minimum norm estimated solution to Equation 2 is *p* = *Mb*.

To correct for superficial sources appearing on the cortical gyri nearest the MEG sensors, we use a depth weighted source covariance matrix which is a diagonal matrix whose diagonals are the norms of the corresponding lead-field matrix column [19],

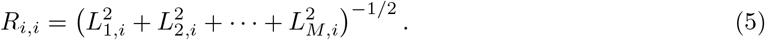

We can further bias our solutions towards the most statistically relevant dipoles using dynamic statistical parametric mapping (dSPM) [2]. The dSPM solution is obtained from normalizing the recon-structed sources by

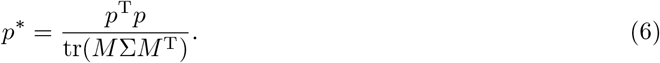

Then *p*^*\**^ is the vector of depth-corrected reconstructed source strengths.

### 2.3 Generation of synthetic data for the evoked N1m peak

We simulated noiseless data for the N1m peak of binaural auditory stimulation. This is similar to the simulation of evoked somatosensory data described in [28], with the difference that to model activation on the primary auditory cortex we used multiple dipoles distributed over a patch of the region rather than a single dipole. We generate this simulated data in two manners, the first using direct BEM-FMM and the second using MNE-Python’s `SourceSimulator` functionality.

To generate the BEM-FMM simulated data, we manually segmented the Heschl gyri of each participant from their T1-weighed MRI data by selecting nearby points in MRI coordinates. The nearest WM mesh triangles were tagged, and a dipole was placed directly above each of the tagged triangle centers between the WM and GM interfaces. The orientations of each of these dipoles were chosen to be the average normal to the nearest GM triangles. Once the dipoles were placed, we used the method of BEM-FMM with *b*-refinement described in [40], to calculate the electric potential on each tissue interface; see also [29]. Once the potential is calculated, we computed the *B*-field on 64 observation points for each magnetometer coil and each individual coil the gradiometer pairs. With this, the integral of the normal component of the *B*-field is computed to generate the sensor outputs of the magnetometers, and in the case of gradiometers we oriented the unit normals in opposite ways for each gradiometer coil and divided the integral by the distance separating the coils of each gradiometer pair, cf. [19]. This process of simulation of the evoked N1m is illustrated in Figure 1.

**Figure 1.**
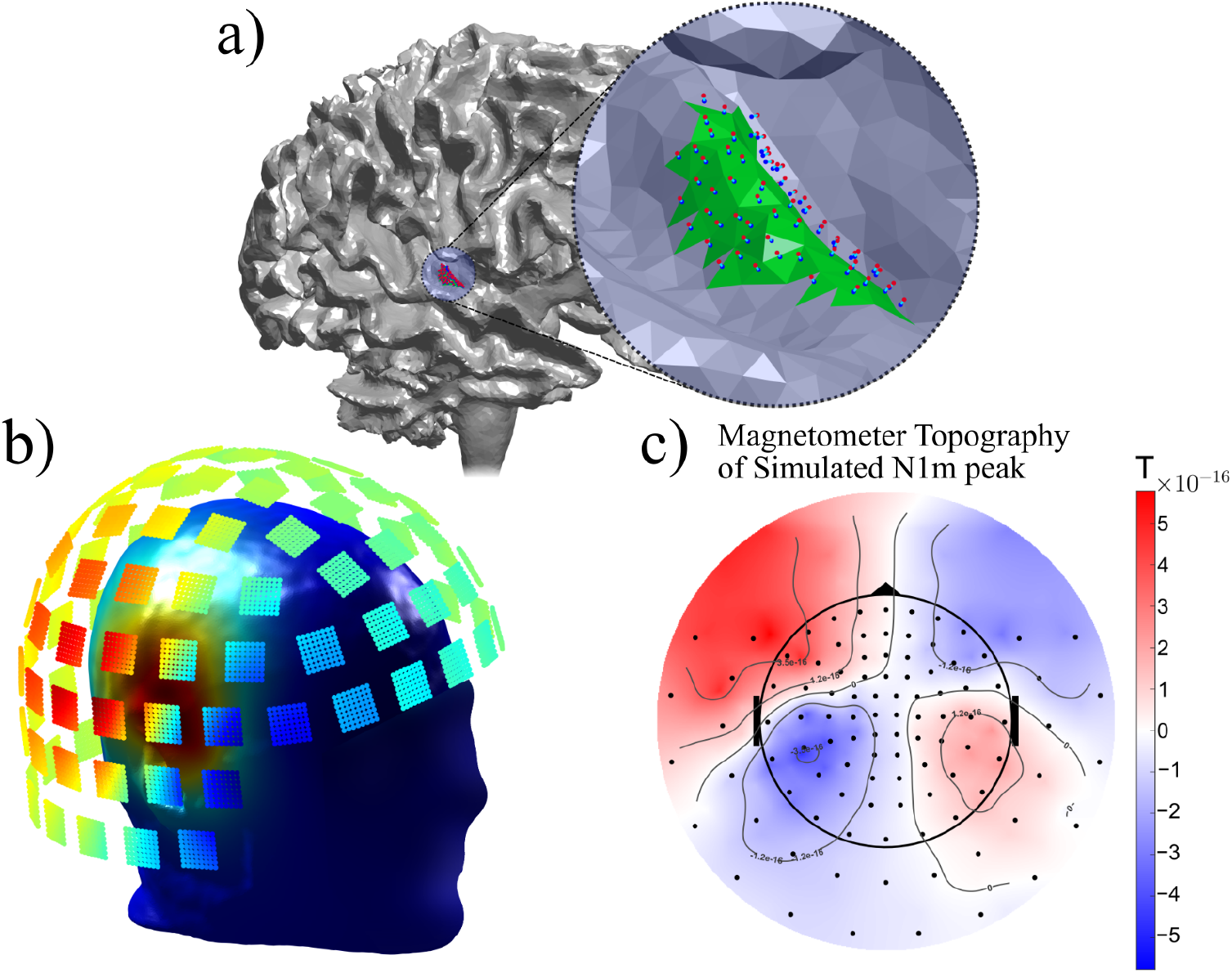
Simulation of the magnetometer signals for the evoked N1m peak on participant N04: a) a patch of WM points (green) at the Heschl gyrus is manually segmented from MRI data; finite-length current dipoles are placed directly above and oriented normal to the cortical surface, with the red and blue spheres denoting the isotropic current sources and sinks, respectively. As such, the direction of each dipole moment is given by the vector from the blue sphere to the red sphere; b) The potential on each tissue (skin plotted) is calculated with BEM-FMM and *b*-refinement, the *B*-field is subsequently computed at 64 observation points per magnetometer coil; c) a surface integral is computed to obtain the sensor outputs; here the interpolated and projected topography of the simulated N1m peak is shown.

The sensor outputs were simulated independently for the dipoles on the Heschl gyri of each hemisphere. The strength of the dipoles was normalized by the number of placed dipoles to make the total strength of the dipoles placed on each hemisphere exactly 1 × 10^−5^ A m. To obtain the sensor outputs for the binaural N1m peak, we simply added the outputs of the activation of the left and right hemisphere — this process is valid by the quasi-static approximation to Maxwell’s equations [34, 27].

For the simulated source outputs using MNE-Python, the regions of interest were chosen according to the FreeSurfer `aparc.a2009s` atlas [3] using the label `G temp supG T transv` across both the left and right hemispheres. From the centroids of these regions, we select all sources within a 10mm extent. These sources are then activated with a sinusoidal waveform time series via MNE-Python’s `mne.simulation.SourceSimulator` protocol [12]. We performed source reconstruction with these simulated sensor outputs at time *t* = 90ms.

### 2.4 Auditory response data processing

#### Data acquisition

Data were provided to us by the authors of [4]. These data were collected from 9 study participants with major depressive disorder. Out of these 9 participants (labeled N01–N09), two of them (N03 and N05) did not have MRI data available, so we restricted our study to the remaining 7. Participants were recruited from the University of New Mexico Mental Health Center to participate in a study focused on the loudness dependence of auditory evoked potentials (LDAEP) in pre-and post-electroconvulsive therapy (ECT). However, for this paper we only analyze the pre-treatment data.

T1-weighted structural MRI images were recorded using the 3-Tesla Siemens Trio MRI scanner at the Mind Research Network of Albuquerque, New Mexico. Raw MEG recordings were collected using the Elekta Neuromag VectorView 306, which possesses 102 magnetometers and 204 gradiometer pairs. Participants were stimulated with binaural auditory tones at a 2kHz frequency, with each tone lasting 50ms. The stimulation was performed using tones of varying loudness; however, for this paper we only consider the 95dB tones. Further details of patient data acquisition are described in [4].

#### Boundary element method models

For the reciprocal BEM-FMM models, the participant MRI images were segmented into seven compartments using the SimNIBS `headreco` pipeline [25] to create the BEM headmodels. BEM models for MNE-Python were segmented using the Freesurfer pipeline [5]. The conductivity values assigned to each BEM model can be seen in Table 2.

**Table 2:** Conductivity values of each tissue layer for the reciprocal BEM-FMM and MNE-Python models.

| Tissue | Conductivity [$\text{S m}^{-1}$] | Reference |
| --- | --- | --- |
| Scalp | 0.25 | [37] |
| Skull | 0.01 | [39] |
| CSF | 1.654 | [39] |
| GM | 0.275 | [39, 36] |
| WM | 0.126 | [39, 36] |
| Ventricles | 1.654 | [39]<sup>†</sup> |
| Eyes | 0.5 | [7, 30] |

(b) Tissue layers in the MNE-Python forward models and their corresponding conductivities and references.
| Tissue | Conductivity [$\text{S m}^{-1}$] | Reference |
| --- | --- | --- |
| Scalp | 0.25 | [37] |
| Outer Skull | 0.01 | [39] |
| Inner Skull | 0.33 | [38] |

#### MEG signal pre-processing

MEG data pre-processing steps for the MPI pipeline were performed in MNE-Python 1.8.0 [11]. Measurement data were run through a FIR low-pass filter with the upper pass-band edge at 85Hz, processed with Maxwell filtering —spatiotemporal signal space separation (tSSS), see [35]— and finally high-pass filtered with a 1Hz lower pass-band edge. Maxwell filtering was done with the default recommended expansion settings of *l*_*in*_ = 8 and *l*_*out*_ = 3, a spatiotemporal buffer duration of 20s, and a spatiotemporal correlation setting of 0.9. The applicability of the default expansion settings was evaluated for a single patient’s data and found adequate. N1m peaks for each participant were determined at a mean latency of 105ms*±*20 ms.

### 2.5 Source reconstruction

In the reciprocal BEM-FMM source reconstruction, individual head-to-MEG coregistration was performed using an iterative closest points (ICP) algorithm [1]. For the MNE-Python source reconstruction pipeline, head-to-MEG coregistration was performed interactively using MNE-C 2.7.3 [12]. All source localization solutions shown in this paper were found using dSPM. For source localization of synthetic N1m peaks, the noise covariance was set to be the identity matrix, with a noise regularization SNR = 100. For localization using the experimental data, we created a noise covariance matrix from the pre-baseline signals at the [− 200, − 5]ms window. The signal-to-noise ratio (SNR) was set at SNR = 1.5 for participants N06 and N08 because their averaged evoked time series were still noisy post-filtering; for all other participants we set SNR = 3. For source reconstruction performed using MNE-Python, we used a depth weighting parameter of 0.65, as this parameter yielded the visually best results. Source reconstruction using reciprocal BEM-FMM was performed with a depth weighting of 1, which is the default parameter setting.

The source space in MNE-Python is found via the `ico-5` option of MNE-Python, which creates a uniform sampling of~ 10242 sources on the interface between the FreeSurfer-segmented WM and GM meshes. On the other hand, the reciprocal BEM-FMM solution uses the triangle centers of the entire SimNIBS `headreco` WM mesh as a source space. To compare the source reconstructions using these two source models, we mapped the MNE-Python source space to the SimNIBS `headreco` mesh using a rotation and translation provided by ICP. For each source (i.e. triangle center) on the `headreco` WM mesh, we computed its source strength by averaging the estimated source strengths of those sources of the MNE-Python source model within a sphere of radius 10mm from the triangle center.

### 2.6 Model analysis using ROC curves

To analyze model performance, we compute the receiver operating characteristic (ROC) curve for each of the source localization solutions using the simulated data. The ROC curve is a useful statistical tool for evaluating the performance of a binary classification model [16]. The ROC curve is computed for source localization by treating the source strengths as binary classification variables across varying activation thresholds [19]. Specifically, we define each of the triangle faces denoting the Heschl gyrus found in Section 2.3 as true dipoles, and all other triangle faces as false dipoles. Given the normalized dSPM solution of source localization, consider a reconstructed source or normalized strength *s* and a threshold *T* from 0 to 1. If the source lies on the tagged Heschl gyrus, we call the source a true positive (TP) if *s* > *T*, a false negative (FN) if *s* < *T*. Similarly, if the source does not lie on the Heschl gyrus, we call the source a false positive (FP) if *s* > *T*, a true negative (TN) if *s* < *T*. The ROC curve is defined as the curve plotting the false positive rate, *FP/*(*FP* + *TN*), versus the true positive rate, *TP/*(*TP* + *FN*), across all thresholds *T*.

The area under the ROC curve (AUC) can be used as a scalar metric to evaluate the performance of the solutions. In particular, if *AUC* = 1, then source localization perfectly captures the true dipoles tagged on the Heschl gyrus. If *AUC* ≤ 0.5, then the source localization is no better than a random chance solution.

The smoothing that is done on the MNE-Python source estimations may negatively bias the ROC curves. Thus, when computing ROC curves for these solutions, we only consider the sparse MNE-Python solution and define the true dipoles as those on the sparse white matter mesh that are within 10mm of the triangle centers of the tagged Heschl gyrus. The TP, FN, FP, and TN sources are then computed in the same manner as described above.

### 2.7 Quantifying solution focality via dispersion

We can quantify the focality of the source estimation solutions by obtaining a measurement of solution dispersion. This is accomplished by constructing a probability density function from the normalized source estimates *s*_*i*_. Specifically, we compute *ŝ*_*i*_ = *αs*_*i*_ where *α* is a constant so that *ŝ*_*i*_ = 1. In this manner, *ŝ* is a probability density function with Cartesian coordinates treated as random variables. We then measure the dispersion of the estimated sources by computing the standard deviation of the distribution across *X, Y*, and *Z* coordinates. A smaller standard deviation, and thus a smaller dispersion, indicates a more focal solution and vice versa. Of course, this measurement depends on anatomical structure, and therefore cannot be used for precise comparison across participants. But we may make use of this to compare the focality of different solutions on the same participant.

Each white matter is aligned with the Cartesian axis, with the lateral, anterior-posterior, and superior-inferior axes corresponding to the *X, Y*, and *Z* axes, respectively. The binaural stimulation of the experimental data is therefore expected to yield symmetric activation across the *X* direction (i.e. across the left and right hemispheres). It follows that the distribution *ŝ* would be bimodal in the *X* direction. In this case, the standard deviation would misrepresent the focality of the source estimates. Because of this, we construct two distributions in the same manner described above, each corresponding to the left and right hemispheres. Focality of the source estimates can then be measured across each hemisphere separately.

## 3 Results

Figure 2 shows source localization results for the simulated N1m peak data on participant N04. Figure 2a shows the solutions found from reciprocal BEM-FMM and MNE-Python using both magnetometer and gradiometer data. Figure 2b shows the sensor topography of the simulated magnetometer and gradiometer data. Figure 2c shows the ROC curves and corresponding AUC values for participant N04. Tables 3a and 3b show the computed AUC values corresponding to each participants source localization solutions from simulated magnetometer and gradiometer data, respectively. The estimated sources and corresponding ROC curves for every participant may be found in the Supplemental Figures.

**Table 3:**
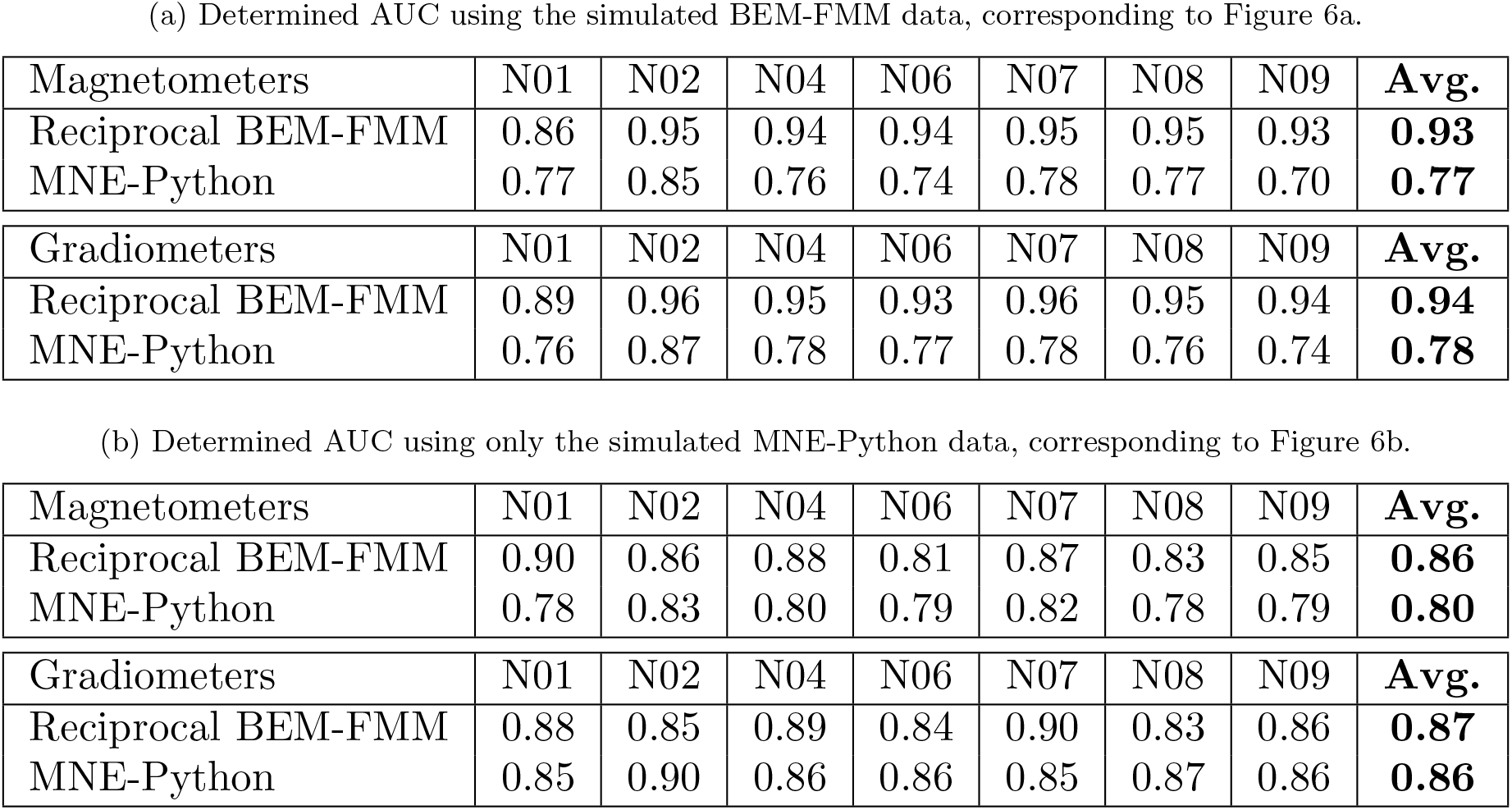
The AUC values of the source localization models at the simulated N1m peak across all participants using reciprocal BEM-FMM and MNE-Python.

|  |  |  |  |  |  |  |  |  |
| --- | --- | --- | --- | --- | --- | --- | --- | --- |
| Magnetometers | N01 | N02 | N04 | N06 | N07 | N08 | N09 | **Avg.** |
| Reciprocal BEM-FMM | 0.86 | 0.95 | 0.94 | 0.94 | 0.95 | 0.95 | 0.93 | **0.93** |
| MNE-Python | 0.77 | 0.85 | 0.76 | 0.74 | 0.78 | 0.77 | 0.70 | **0.77** |
| Gradiometers | N01 | N02 | N04 | N06 | N07 | N08 | N09 | **Avg.** |
| Reciprocal BEM-FMM | 0.89 | 0.96 | 0.95 | 0.93 | 0.96 | 0.95 | 0.94 | **0.94** |
| MNE-Python | 0.76 | 0.87 | 0.78 | 0.77 | 0.78 | 0.76 | 0.74 | **0.78** |
| (b) Determined AUC using only the simulated MNE-Python data, corresponding to Figure 6b. |  |  |  |  |  |  |  |  |
| Magnetometers | N01 | N02 | N04 | N06 | N07 | N08 | N09 | **Avg.** |
| Reciprocal BEM-FMM | 0.90 | 0.86 | 0.88 | 0.81 | 0.87 | 0.83 | 0.85 | **0.86** |
| MNE-Python | 0.78 | 0.83 | 0.80 | 0.79 | 0.82 | 0.78 | 0.79 | **0.80** |
| Gradiometers | N01 | N02 | N04 | N06 | N07 | N08 | N09 | **Avg.** |
| Reciprocal BEM-FMM | 0.88 | 0.85 | 0.89 | 0.84 | 0.90 | 0.83 | 0.86 | **0.87** |
| MNE-Python | 0.85 | 0.90 | 0.86 | 0.86 | 0.85 | 0.87 | 0.86 | **0.86** |

**Figure 2.**
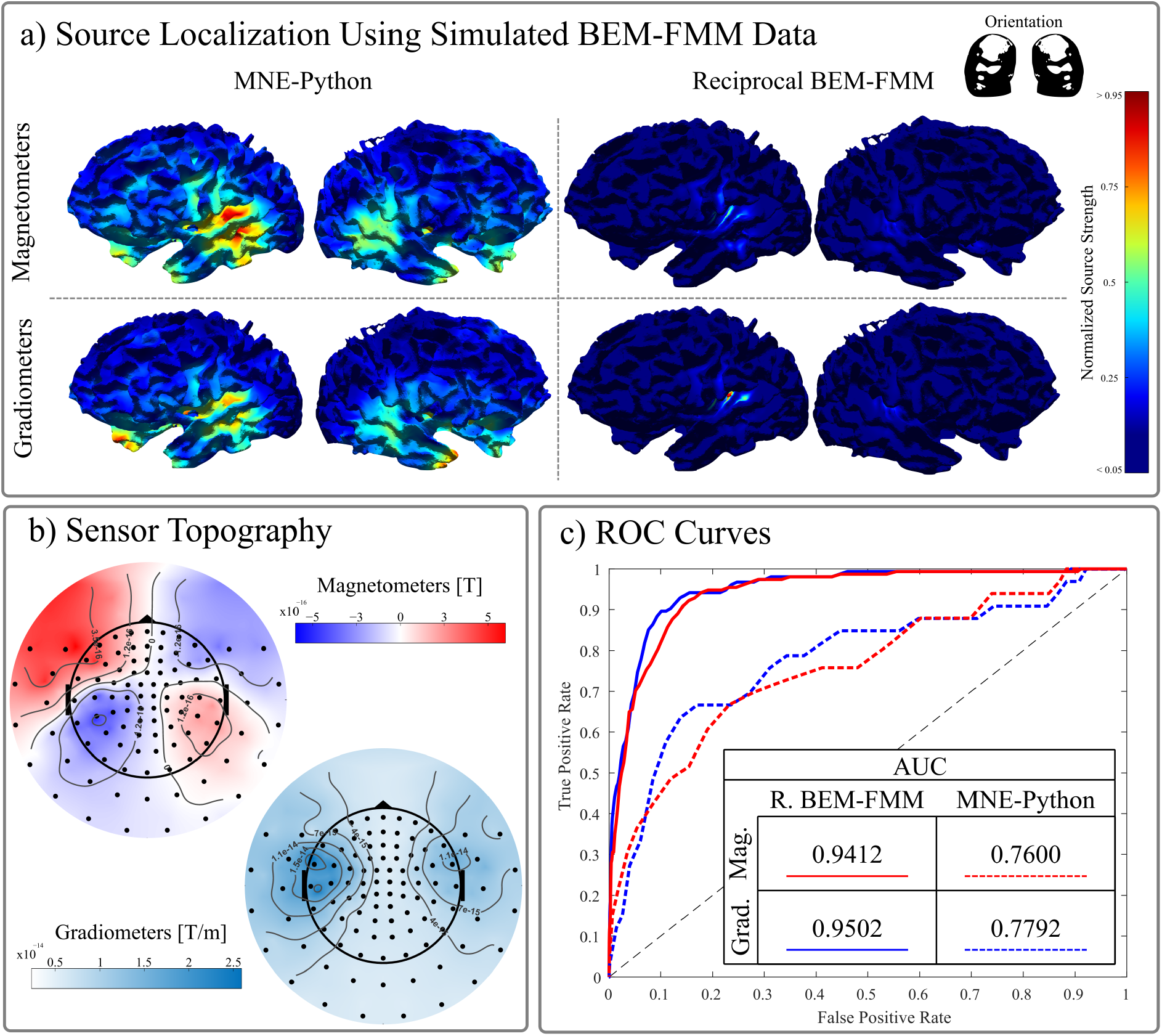
Source localization and ROC curve analysis for participant N04 using synthetic data. a) The source localization solutions using the simulated N1m peak data using both reciprocal BEM-FMM and MNE-Python in the left and right columns, respectively. The top row indicates solutions found using magnetometer data, and the bottom row that of gradiometer data. Perspectives are shifted to achieve a better view of the Heschl gyri. Peak activation for both sensor types appears on the posterior-superior part of the left Heschl gyrus for both source localization methods, with reciprocal BEM-FMM achieving very focal activation. b) The sensor topography of the simulated N1m peak data is shown for both magnetometers and gradiometers. c) The ROC curves of each source reconstruction are shown, with reciprocal BEM-FMM in solid lines and MNE-Python in dashed lines. The red lines indicate solutions from magnetometer data, and the blue lines that of gradiometer data. The table of values denotes the AUC for each of the corresponding models.

Figure 3 shows the source reconstruction results for the measured N1m peak data of participant N01 using both MNE-Python and reciprocal BEM-FMM, with Figure 3a showing the evoked gradiometer signals and topography, and Figure 3b showing the dSPM solutions using each of the two solvers. Figure 4 shows detailed source reconstruction results for participant N09 using the magnetometer data, with Figure 4a and Figure 4b showing the normalized source strengths on the left and right white matter hemispheres, respectively, and Figure 4c showing the corresponding probability distributions and focality measurements. Figure 5 and Figure 6 show the dSPM source reconstructions for all participants using MNE-Python and reciprocal BEM-FMM, with Figure 5 showing the reconstructions using the N1m magnetometer peaks, and Figure 6 showing the reconstructions using the N1m gradiometer peaks.

**Figure 3.**
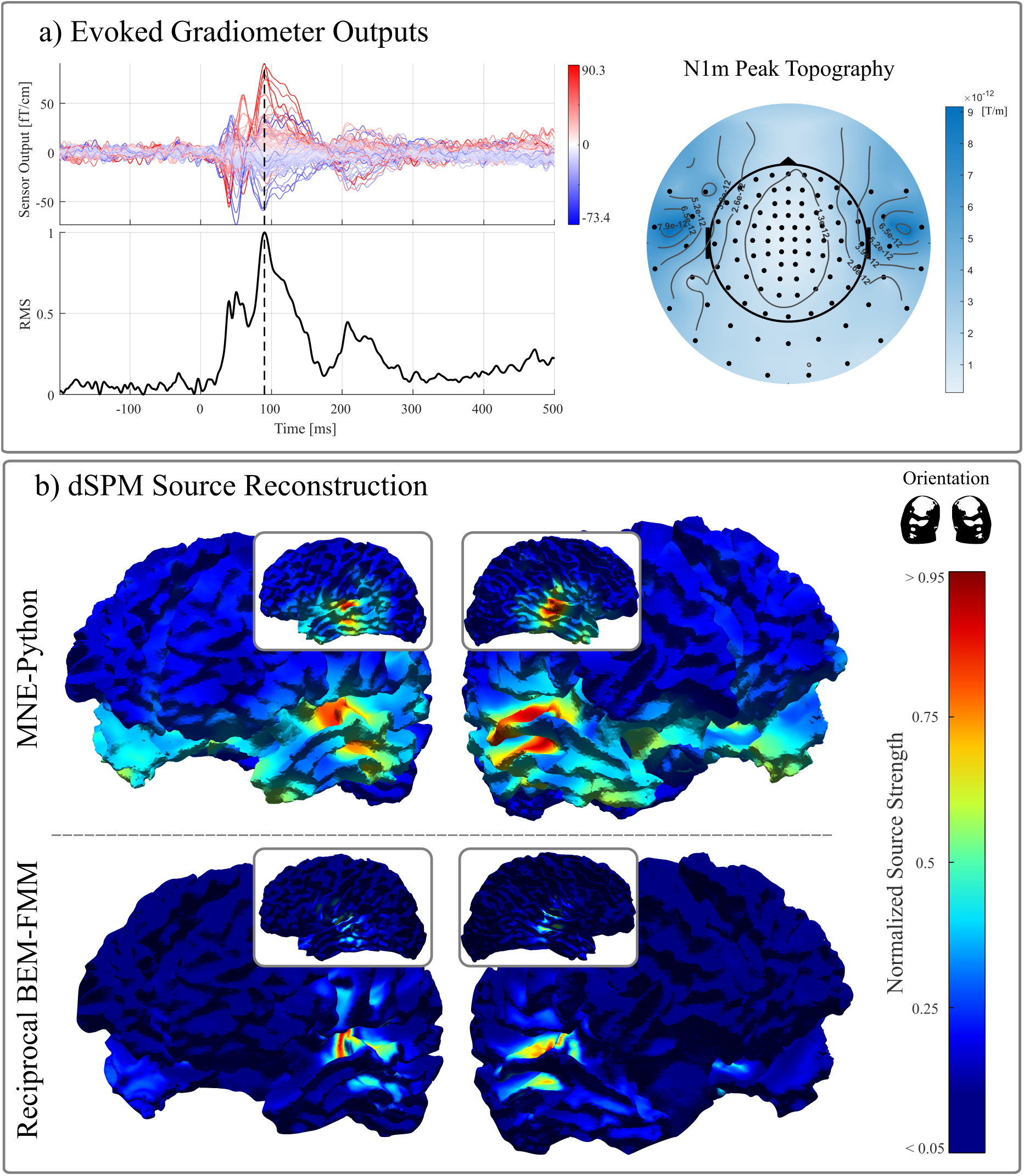
Source reconstruction results of participant N01 using the gradiometer data with both MNE-Python and reciprocal BEM-FMM. a) The evoked gradiometer sensor data is shown on the left, along with the signal RMS. The N1m peak is marked with the dotted black line, and the sensor topography at this peak is shown on the right. b) The normalized dSPM solutions are shown, with the smoothed MNE-Python solution shown on the top, and the reciprocal BEM-FMM solution shown on bottom. The 18 left and right columns show focus perspectives of the left and right hemispheres, respectively. The inset images show the solution on each hemisphere from a lateral perspective.

**Figure 4.**
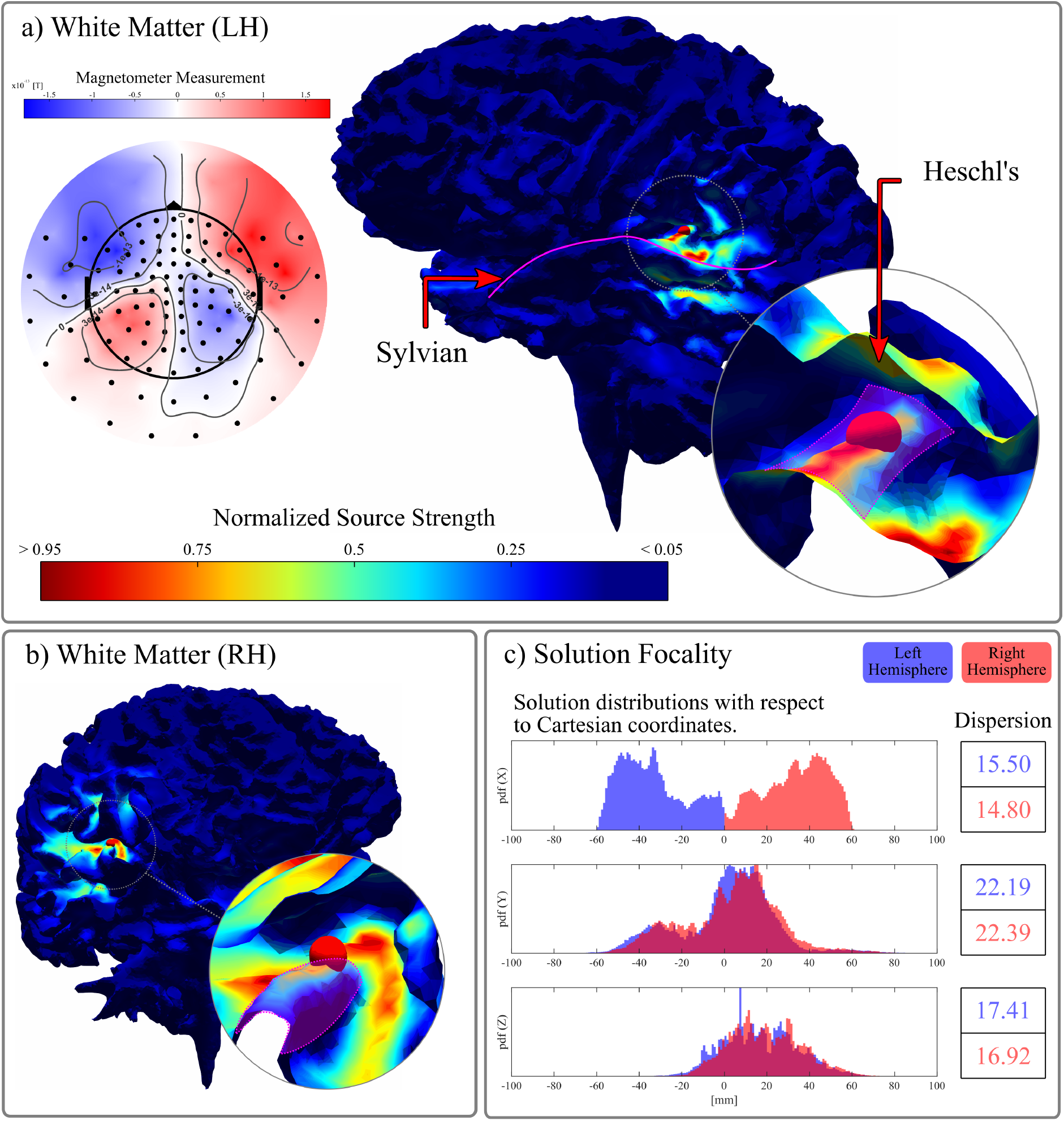
Detailed source reconstruction results for participant N09 using the magnetometer sensor data with reciprocal BEM-FMM. a) To the left, the magnetometer topography at the N1m peak is shown. To the right, the reconstructed source strengths are plotted on the white matter surface, with visual focus emphasizing the left hemisphere. The Sylvian fissure is outlined with the magenta curve. The source of strongest strength (activation peak) is denoted as the red sphere. In the magnified view, a portion of the Heschl gyrus is highlighted in magenta. b) The same reconstructed source strengths are shown, with visual focus now on the right hemisphere, and the Heschl gyrus highlighted again in magenta. c) The source estimate distributions along the Cartesian coordinates. The distribution from the left and right hemispheres are shown in blue and pink, resp1e9ctively. The corresponding dispersion measures for each distribution are shown on the right.

**Figure 5.**
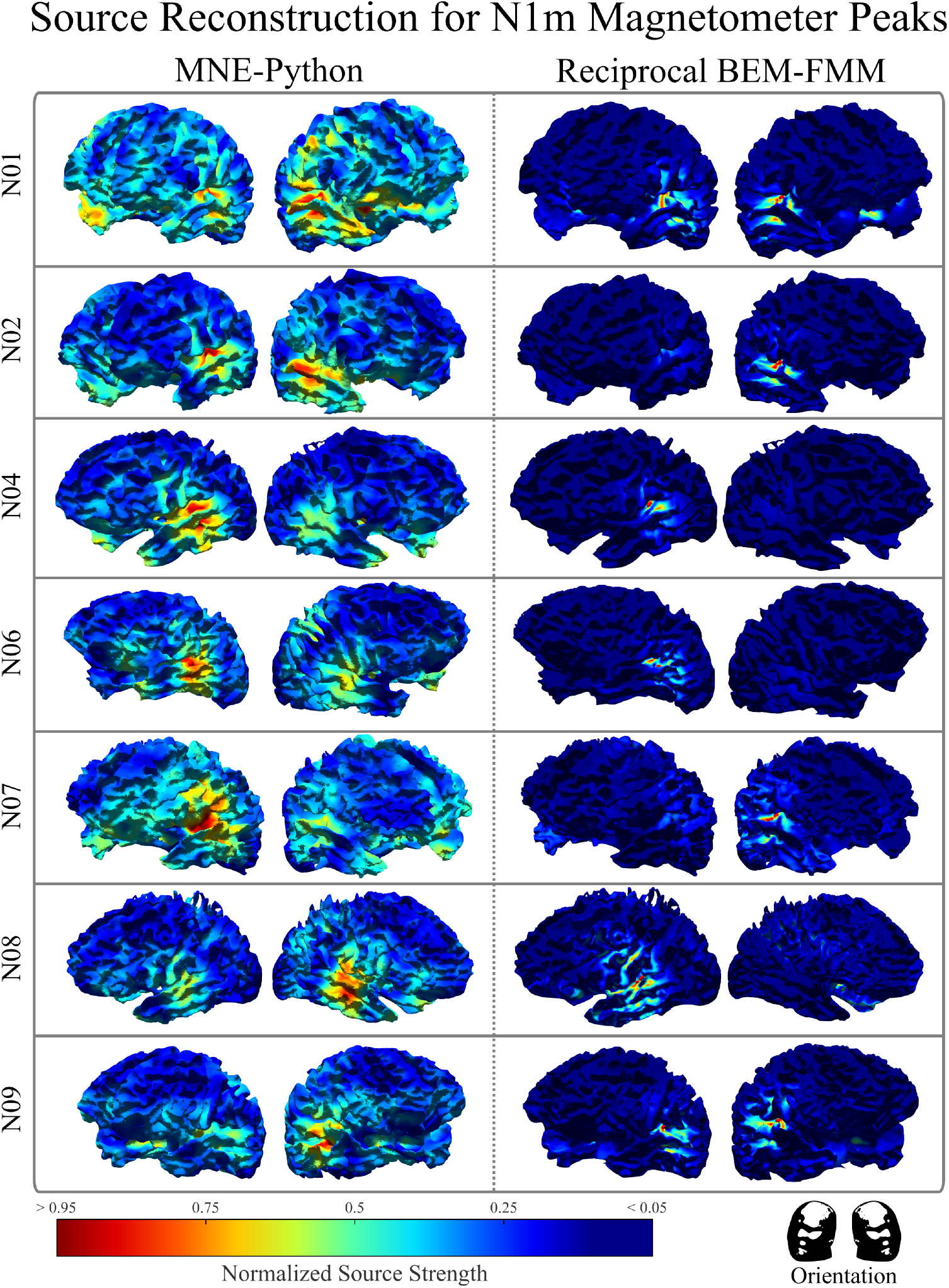
Source localization results of all participants at their respective auditory evoked N1m magnetometer peaks using reciprocal BEM-FMM and MNE-Python. For each participant, white matter perspectives are chosen to focus on the temporal lobe and Heschl gyrus in the top and bottom rows, respectively, on both the left and right hemispheres. The color map indicates the normalized current density, with blue and red colors indicating 0 and 1 values, respectively.

**Figure 6.**
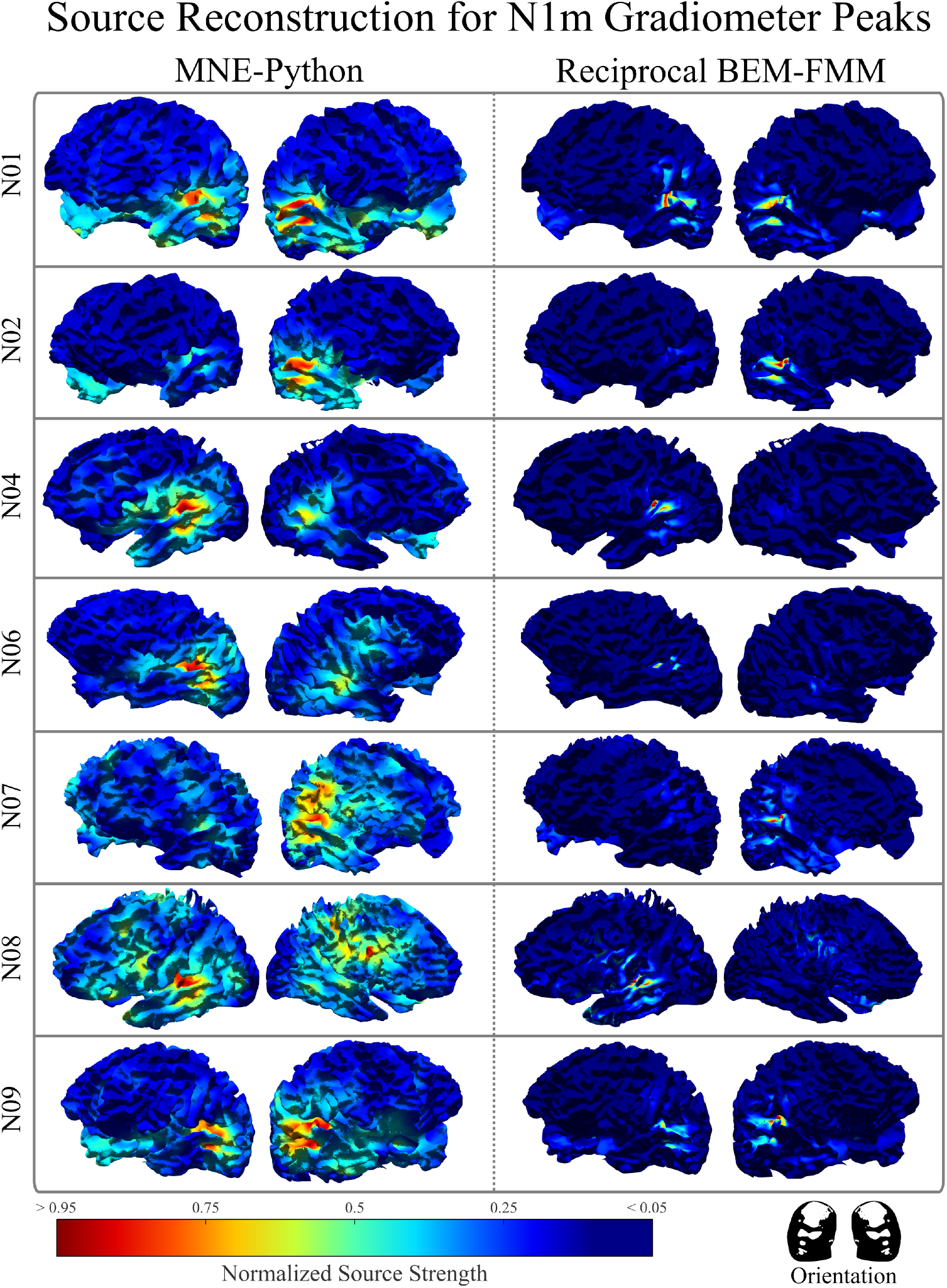
Source localization results of all participants at their respective auditory evoked N1m gradiometer peaks using reciprocal BEM-FMM and MNE-Python. For each participant, white matter perspectives are chosen to focus on the temporal lobe and Heschl gyrus in the top and bottom rows, respectively, on both the left and right hemispheres. The color map indicates the normalized current density, with blue and red colors indicating 0 and 1 values, respectively.

Tables 4a and 4b show the measured dispersion of the source localizations for every participant using MNE-Python and reciprocal BEM-FMM. From these values, the corresponding percent improvements from MNE-Python to reciprocal BEM-FMM are shown in Tables 5a and 5b. Additional figures showing the probability density functions for every subject may be found in the Supplemental Figures.

**Table 4:** Dispersion measurements of the estimated sources of every participant using MNE-Python and reciprocal BEM-FMM. The dispersion is measured by treating the normalized source estimates as a probability density function and computing the standard deviation. Dispersion measurements are separated by the left and right hemisphere. The standard deviation is shown for each of the Cartesian coordinates, and the average of the three standard deviations is computed.

| Dispersion (Magnetometers) |  |  |  |  |  |  |  |  |
| --- | --- | --- | --- | --- | --- | --- | --- | --- |
| MNE | Left Hemisphere |  |  |  | Right Hemisphere |  |  |  |
| Subject | X | Y | Z | Avg. | X | Y | Z | Avg. |
| *N01* | 16.79 | 33.20 | 24.71 | 24.90 | 17.80 | 31.07 | 24.30 | 24.39 |
| *N02* | 17.10 | 31.39 | 20.07 | 22.86 | 17.41 | 31.53 | 20.56 | 23.17 |
| *N04* | 17.35 | 31.93 | 20.95 | 23.41 | 18.07 | 33.03 | 20.86 | 23.99 |
| *N06* | 16.12 | 31.42 | 20.52 | 22.68 | 16.18 | 31.88 | 22.09 | 23.38 |
| *N07* | 15.27 | 30.46 | 21.50 | 22.41 | 16.82 | 29.13 | 21.71 | 22.56 |
| *N08* | 18.59 | 33.74 | 23.29 | 25.21 | 17.25 | 32.56 | 22.62 | 24.14 |
| *N09* | 15.36 | 32.74 | 21.84 | 23.31 | 17.02 | 31.27 | 21.54 | 23.27 |

| R.BEM-FMM | Left Hemisphere |  |  |  | Right Hemisphere |  |  |  |
| --- | --- | --- | --- | --- | --- | --- | --- | --- |
| Subject | X | Y | Z | Avg. | X | Y | Z | Avg. |
| *N01* | 14.06 | 19.73 | 16.99 | 16.93 | 17.41 | 21.38 | 18.02 | 18.94 |
| *N02* | 14.72 | 21.25 | 16.42 | 17.46 | 15.29 | 18.94 | 15.99 | 16.74 |
| *N04* | 15.79 | 22.81 | 18.37 | 18.99 | 15.62 | 21.33 | 17.04 | 18.00 |
| *N06* | 15.39 | 23.86 | 22.24 | 20.50 | 15.09 | 21.00 | 19.25 | 18.44 |
| *N07* | 14.64 | 24.19 | 20.57 | 19.80 | 12.53 | 18.48 | 16.39 | 15.80 |
| *N08* | 18.94 | 26.12 | 20.35 | 21.80 | 14.97 | 21.44 | 18.02 | 18.15 |
| *N09* | 15.50 | 22.19 | 17.41 | 18.36 | 14.80 | 22.39 | 16.92 | 18.04 |

(b) Dispersion measurements from solutions computed using N1m gradiometer peaks.
| Dispersion (Gradiometers) |  |  |  |  |  |  |  |  |
| --- | --- | --- | --- | --- | --- | --- | --- | --- |
| MNE | Left Hemisphere |  |  |  | Right Hemisphere |  |  |  |
| Subject | X | Y | Z | Avg. | X | Y | Z | Avg. |
| *N01* | 16.69 | 32.61 | 23.73 | 24.34 | 17.64 | 31.05 | 23.87 | 24.19 |
| *N02* | 17.04 | 28.54 | 19.81 | 21.80 | 17.42 | 28.08 | 20.08 | 21.86 |
| *N04* | 17.37 | 30.29 | 19.75 | 22.47 | 17.67 | 30.47 | 20.15 | 22.76 |
| *N06* | 16.38 | 29.98 | 19.40 | 21.92 | 16.33 | 29.86 | 21.73 | 22.64 |
| *N07* | 15.29 | 30.05 | 22.32 | 22.55 | 17.10 | 29.89 | 22.27 | 23.08 |
| *N08* | 18.61 | 32.17 | 21.96 | 24.25 | 17.08 | 31.36 | 20.91 | 23.11 |
| *N09* | 15.54 | 32.31 | 19.63 | 22.49 | 17.24 | 30.10 | 20.23 | 22.52 |

| R.BEM-FMM | Left Hemisphere |  |  |  | Right Hemisphere |  |  |  |
| --- | --- | --- | --- | --- | --- | --- | --- | --- |
| Subject | X | Y | Z | Avg. | X | Y | Z | Avg. |
| *N01* | 13.38 | 18.54 | 14.81 | 15.58 | 16.28 | 20.09 | 16.29 | 17.55 |
| *N02* | 16.17 | 24.38 | 17.00 | 19.18 | 17.01 | 22.14 | 15.53 | 18.23 |
| *N04* | 16.98 | 23.33 | 19.09 | 19.80 | 16.86 | 22.00 | 17.20 | 18.69 |
| *N06* | 16.51 | 28.67 | 19.64 | 21.61 | 15.63 | 24.96 | 17.75 | 19.45 |
| *N07* | 15.74 | 24.95 | 17.51 | 19.40 | 13.14 | 19.89 | 15.67 | 16.23 |
| *N08* | 18.99 | 30.08 | 17.97 | 22.35 | 15.65 | 23.83 | 16.62 | 18.70 |
| *N09* | 16.76 | 24.75 | 16.67 | 19.39 | 15.77 | 24.22 | 16.54 | 18.84 |

**Table 5:** The percent improvement of dispersion of the estimated sources from MNE-Python to reciprocal BEM-FMM, corresponding to the values from Table 4.

| Subject | Percent Improvement (Magnetometers) |  |  |  |  |  |  |  |
| --- | --- | --- | --- | --- | --- | --- | --- | --- |
|  | Left Hemisphere |  |  |  | Right Hemisphere |  |  |  |
|  | X | Y | Z | Avg. | X | Y | Z | Avg. |
| *N01* | 16.2% | 40.6% | 31.2% | 32.0% | 2.2% | 31.2% | 25.8% | 22.4% |
| *N02* | 13.9% | 32.3% | 18.2% | 23.6% | 12.2% | 39.9% | 22.2% | 27.7% |
| *N04* | 9.0% | 28.6% | 12.3% | 18.9% | 13.6% | 35.4% | 18.3% | 25.0% |
| *N06* | 4.5% | 24.0% | -8.4% | 9.6% | 6.7% | 34.1% | 12.9% | 21.1% |
| *N07* | 4.1% | 20.6% | 4.3% | 11.6% | 25.5% | 36.5% | 24.5% | 29.9% |
| *N08* | -1.9% | 22.6% | 12.6% | 13.5% | 13.2% | 34.1% | 20.4% | 24.8% |
| *N09* | -0.9% | 32.2% | 20.3% | 21.2% | 13.0% | 28.4% | 21.4% | 22.5% |

(b) Percent improvement of dispersion from Table 4b.
| Subject | Percent Improvement (Gradiometers) |  |  |  |  |  |  |  |
| --- | --- | --- | --- | --- | --- | --- | --- | --- |
|  | Left Hemisphere |  |  |  | Right Hemisphere |  |  |  |
|  | X | Y | Z | Avg. | X | Y | Z | Avg. |
| *N01* | 19.8% | 43.1% | 37.6% | 36.0% | 7.7% | 35.3% | 31.8% | 27.4% |
| *N02* | 5.1% | 14.6% | 14.2% | 12.0% | 2.4% | 21.1% | 22.7% | 16.6% |
| *N04* | 2.2% | 23.0% | 3.3% | 11.9% | 4.6% | 27.8% | 14.6% | 17.9% |
| *N06* | -0.8% | 4.4% | -1.2% | 1.4% | 4.3% | 16.4% | 18.3% | 14.1% |
| *N07* | -3.0% | 17.0% | 21.5% | 14.0% | 23.2% | 33.4% | 29.6% | 29.7% |
| *N08* | -2.0% | 6.5% | 18.2% | 7.8% | 8.4% | 24.0% | 20.5% | 19.1% |
| *N09* | -7.8% | 23.4% | 15.1% | 13.8% | 8.6% | 19.5% | 18.2% | 16.3% |

## 4 Discussion

### Source estimation of simulated data

The noiseless simulated data (both using 3-layer BEM using MNE-Python’s forward solver, and for 5-layer BEM using direct BEM-FMM with *b*-refinement) was recovered significantly more focally and accurately with our method or reciprocal BEM-FMM (see Figure 2, as well as Figures 2 − 5 in the Supplemental Figures): the estimated sources are distributed almost exclusively along the Heschl gyri, with peaks occuring on one or both of these gyri —an effect that is mostly explained by the positioning of the participant with respect to the MEG sensors. The source reconstructions using MNE-Python show instead spurious regions of activation in many cortical regions away from the Heschl gyri, and a much less focal distribution; nevertheless the peaks remain located near the Heschl gyri with an identical pattern of peak asymmetry observed in the reciprocal BEM-FMM reconstructions.

These conclusions can be also obtained with the statistical analyses of AUC statistics in Table 3. These show that when more realistic forward simulated data is involved (5-layer direct BEM-FMM models), the high-resolution reciprocal BEM-FMM significantly outperforms MNE-Python. When the simplified models are used for the simulated data (3-layer BEM models of MNE-Python), the performance of the reciprocal BEM-FMM is still superior on average, although just marginally —it should be noted that MNE-Python does not recover the sources simulated with the 3-layer BEM perfectly because we activated a cluster of dipoles rather than a single dipole on each of the Heschl gyri.

Given that real experimental data will be more similar to the high-resolution forward models, this indicates that the reciprocal BEM-FMM method will have a better performance than MNE-Python on experimental data. This is confirmed in the following subsection.

### Source estimation of experimental data

The estimated sources recovered by reciprocal BEM-FMM seen in Figure 5 and Figure 6 show very focal activation peaks on the Heschl gyri across all participants, as expected from a binaural auditory evoked N1m peak. In comparison, the estimated sources recovered by MNE-Python achieved activation in the Heschl gyri across all participants. Howver, these regions of activation were much less focalized than that seen in reciprocal BEM-FMM. Further, the MNE-Python solutions possessed spurious regions of activation, which are particularly pronounced in participants N07 and N08.

These focality observations are justified by the dispersion measurements from Tables 4a and 4b, and their corresponding percent improvements in Tables 5a and 5b. These results show that the dispersion of reciprocal BEM-FMM solutions are generally smaller than that of the MNE-Python solutions across all subjects, coordinate axes, and hemispheres. Further, these observations hold true for both magnetometer and gradiometer peaks. For instance, we see that for participant N01, the reciprocal BEM-FMM solutions improve average focality for both sensor types by over 30% on the left hemisphere, and over 20% on the right hemisphere. Similar improvements in focality are seen for the rest of the subjects, with stronger focality improvements for solutions from magnetometer data over those from gradiometer data. Participant N06 appears as an exception to these trends, with only a marginal improvement in focality of 1.4% on the left hemisphere from gradiometer data. However, this is counterbalanced by the substantive focality improvement of 14% on the right hemisphere. That is, even in exceptional cases, reciprocal BEM-FMM yields solutions with significantly less dispersion than those obtained from MNE-Python.

It is worth reiterating that, aside from the SNR values of participants N06 and N08, there was no subject-specific tweaking performed in either reciprocal BEM-FMM or MNE-Python. Both source reconstruction solvers were run, for all intents and purposes, utilizing their default settings. The only major parameter change was in the depth weighting constant, which was chosen to be visually optimal for MNE-Python. It is possible that improvements to the solutions achieved by MNE-Python could be made through additional parameter modifications. However, such subject-specific parameter tweaking may be unnecessary in reciprocal BEM-FMM based on the results found here.

In the comparative source reconstruction results of [28] between our methods and MNE-Python, our methods performed only marginally better. However, for the auditory data that we considered here, our method provides a significant improvement. A possible explanation for this is that the geometry of the head near the central sulcus can be sufficiently approximated by a sphere or by a simplified classical BEM model with low-resolution shells. On the other hand, the temporal lobes cannot be accurately described with simpler models. Another possible explanation for this is that the quality of the evoked data in [28] allowed for simpler low resolution models to perform well. Conversely, the experimental auditory data used here was less than ideal — it posed a fair challenge to process and, even with sufficient filtering, it still seemed subject to moderate noise. With this in mind, we find the highly focal source localizations obtained from reciprocal BEM-FMM to be quite impressive.

## 5 Conclusions

Our results indicate that high-resolution forward models including cortical layers significantly improve the accuracy of source estimates from AEFs compared to that of the 3-layer BEM models in MNE-Python. All of our source estimations using high-resolution models indicate good quality of source localizations, with focal peaks of activation found on the Heschl gyri of each participant. This performance held true across various types of data, both simulated and experimental. Using simulated AEFs and ROC measures, we find statistically superior results to the 3-layer BEM of MNE-Python.

Specifically, source estimates yield an ROC measure of about 80% from MNE-Python, and about 90% from reciprocal BEM-FMM. Results from experimental data showed visually apparent improvement of reciprocal BEM-FMM over the 3-layer BEM. These observations were justified by quantifying the solution focality, where we see reciprocal BEM-FMM solutions possess upwards of 30% reduction in dispersion over MNE-Python. The high resolution source space and focalized source estimations achieved by reciprocal BEM-FMM indicates that, through some further numerical and statistical analysis, our methodology could improve the applications of source localization in clinical settings.

