## Supplemental Figures for "Improved Source Localization of Auditory Evoked Fields using Reciprocal BEM-FMM"

[†]First two authors share primary authorship; last two authors share senior authorship.
[1]Dept. of Electrical & Computer Engineering, Worcester Polytechnic Institute, Worcester, MA, USA
[2]Graduate School of Information Sciences, Division of Mathematics, Tohoku University, Sendai, Miyagi, Japan
[3]Technische Universität Ilmenau, Ilmenau, Thuringia, Germany
[4]Max Planck Institute for Human Cognitive and Brain Sciences, Leipzig, Saxony, Germany
[5]The Mind Research Network, University of New Mexico, Albuquerque, NM, USA
[6]Department of Psychiatry and Behavioral Sciences, University of New Mexico School of Medicine, Albuquerque, NM, USA
[7]Athinoula A. Martinos Center for Biomedical Imaging, Massachusetts General Hospital, Boston, MA, USA
[8]Computational Neurostimulation Research Program, Noninvasive Neuromodulation Unit, Experimental Therapeutics and Pathophysiology Branch, National Institute of Mental Health, National Institutes of Health, Bethesda, MD, USA

July 2025

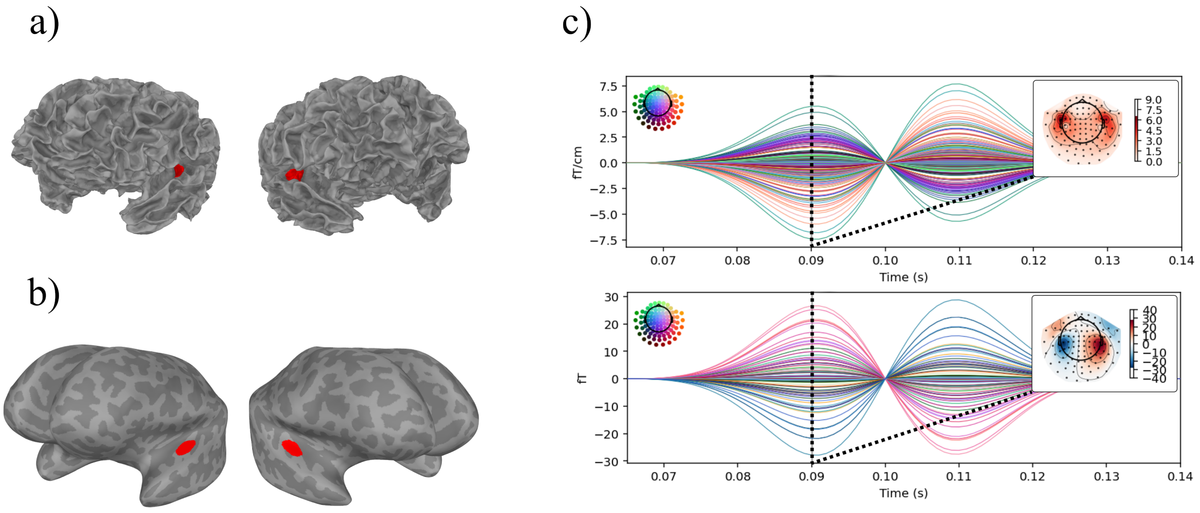


Figure 1: Simulation of sensor outputs for participant N04 using MNE-Python: a) The activated sources on the FreeSurfer white matter mesh. b) The activated sources on the inflated white matter. c) The simulated gradiometer (top) and magnetometer (bottom) sensor outputs according to the activated sources and a sinusoidal waveform, with topographies selected at the time $t = 0.09$. The waveform is constructed with oscillating frequency 15Hz, zero-measure latency 0.1ms.

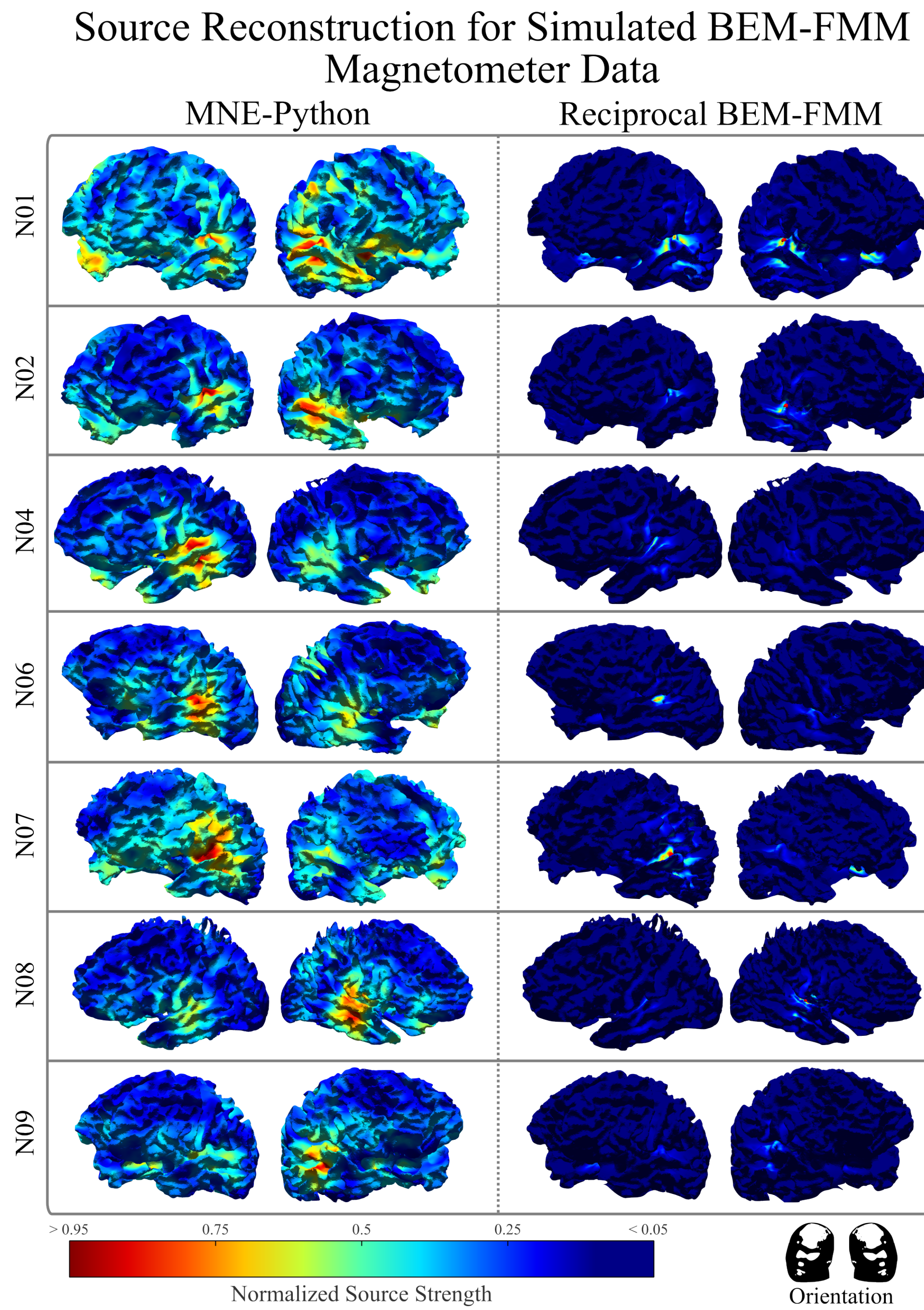


Figure 2: Source localization results of all participants at their respective simulated magnetometer data using reciprocal BEM-FMM and MNE-Python, with the simulated data generated using BEM-FMM. For each participant, white matter perspectives are chosen to focus on the temporal lobe and Heschl gyrus in the top and bottom rows, respectively, on both the left and right hemispheres. The color map indicates the normalized current density, with blue and red colors indicating 0 and 1 values, respectively.

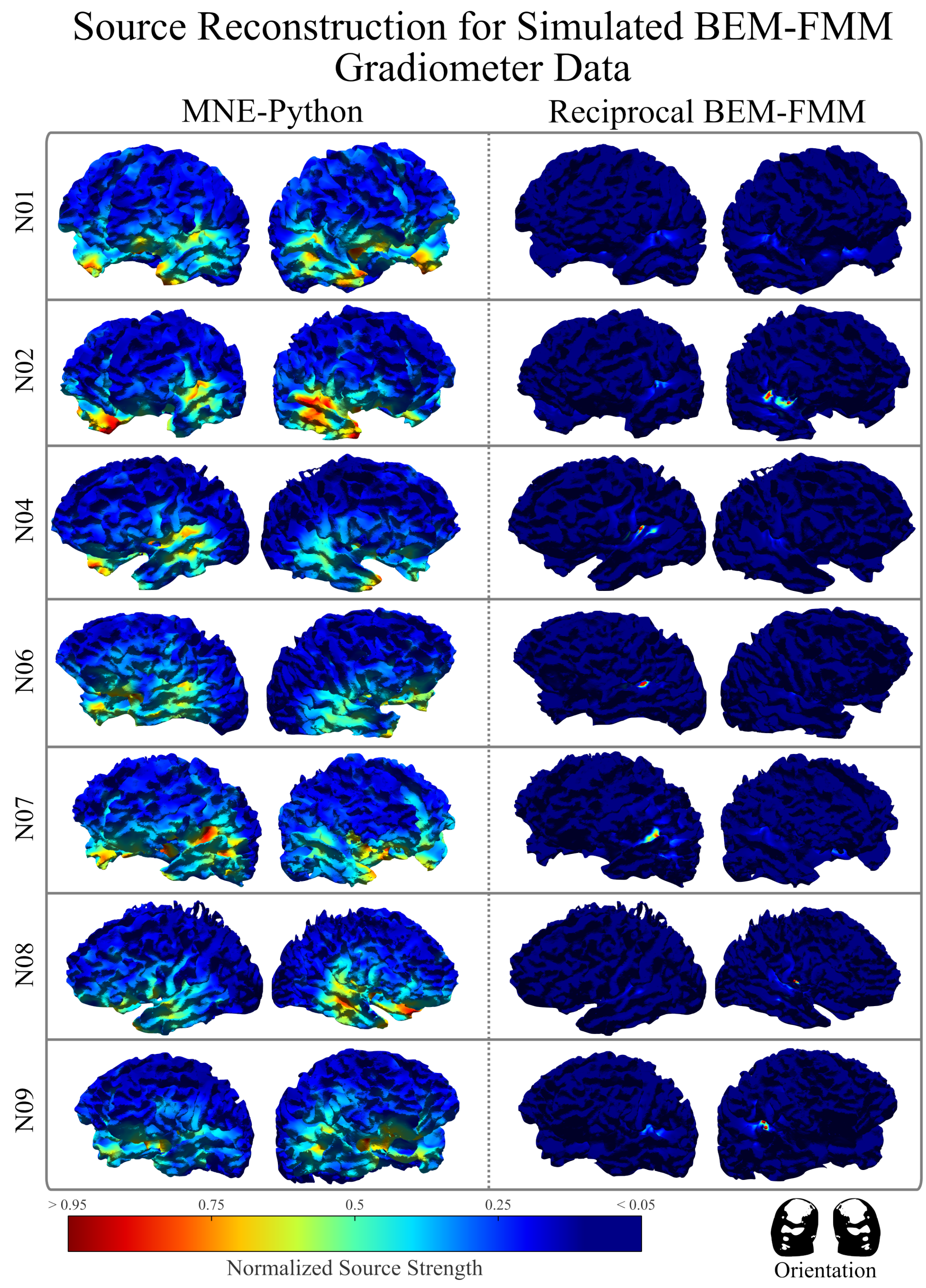


Figure 3: Source localization results of all participants at their respective simulated gradiometer data using reciprocal BEM-FMM and MNE-Python, with the simulated data generated using BEM-FMM. For each participant, white matter perspectives are chosen to focus on the temporal lobe and Heschl gyrus in the top and bottom rows, respectively, on both the left and right hemispheres. The color map indicates the normalized current density, with blue and red colors indicating 0 and 1 values, respectively.

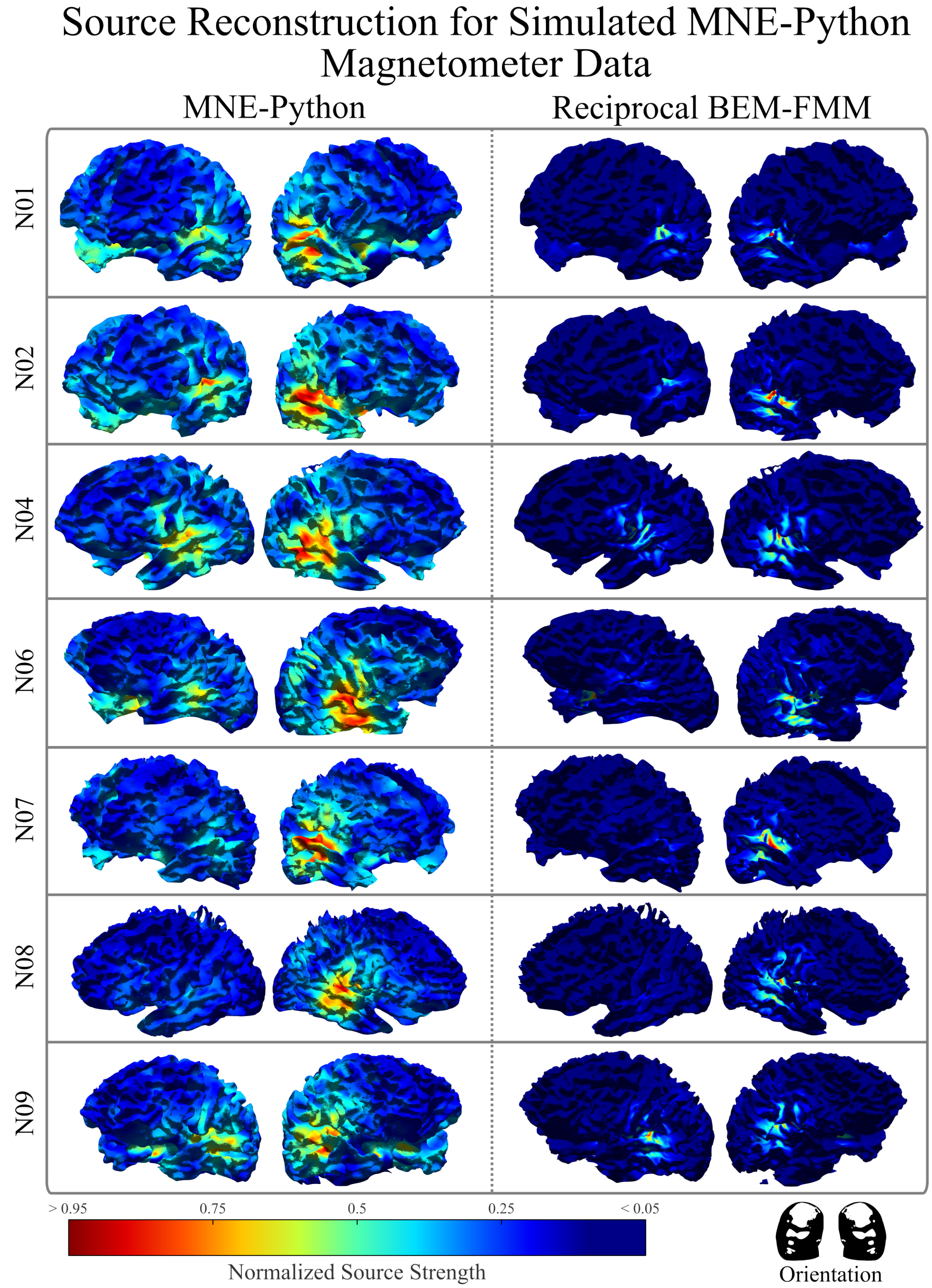


Figure 4: Source localization results of all participants at their respective simulated magnetometer data using reciprocal BEM-FMM and MNE-Python, with the simulated data generated using MNE-Python. For each participant, white matter perspectives are chosen to focus on the temporal lobe and Heschl gyrus in the top and bottom rows, respectively, on both the left and right hemispheres. The color map indicates the normalized current density, with blue and red colors indicating 0 and 1 values, respectively.

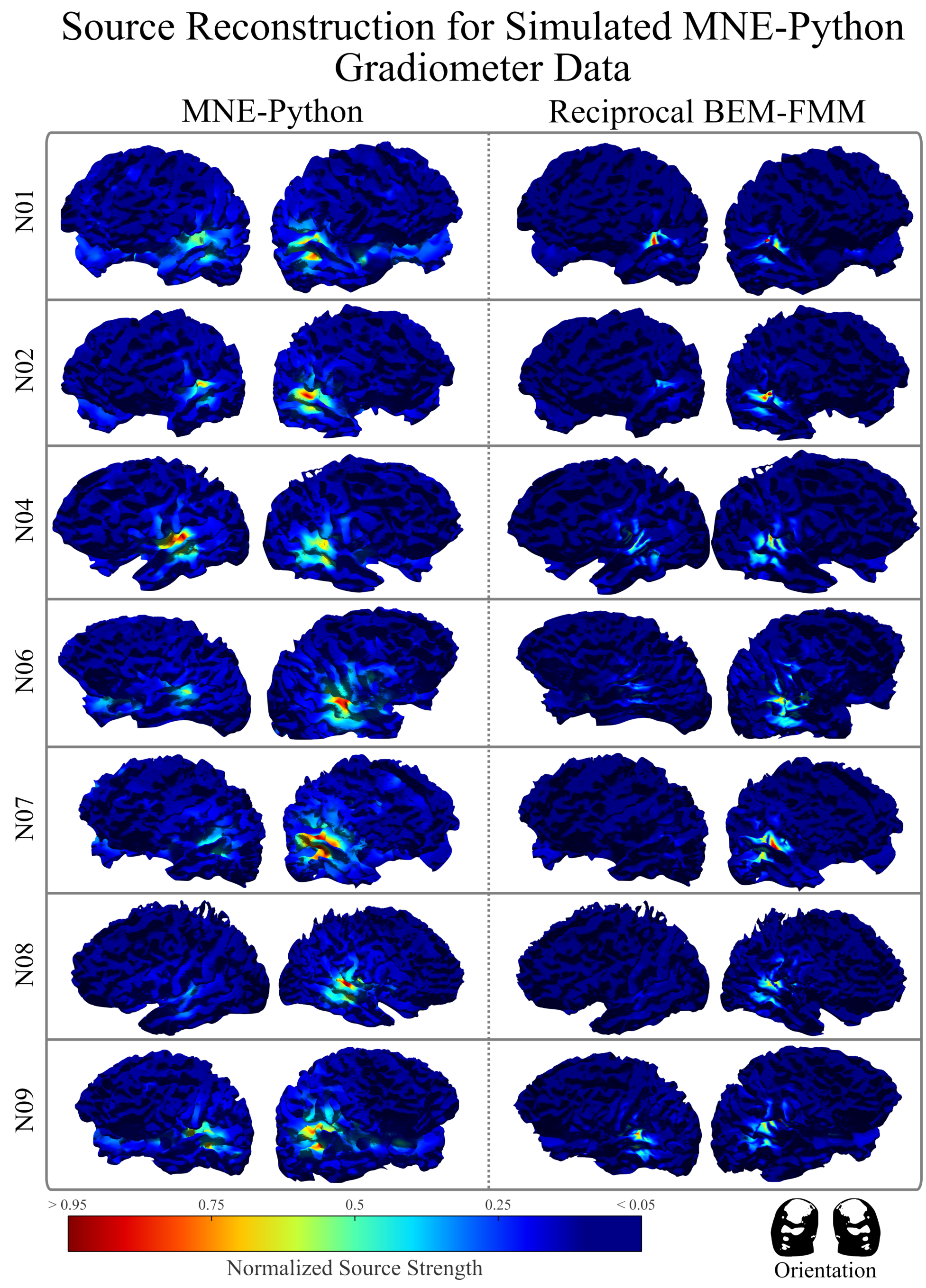


Figure 5: Source localization results of all participants at their respective simulated gradiometer data using reciprocal BEM-FMM and MNE-Python, with the simulated data generated using MNE-Python. For each participant, white matter perspectives are chosen to focus on the temporal lobe and Heschl gyrus in the top and bottom rows, respectively, on both the left and right hemispheres. The color map indicates the normalized current density, with blue and red colors indicating 0 and 1 values, respectively.

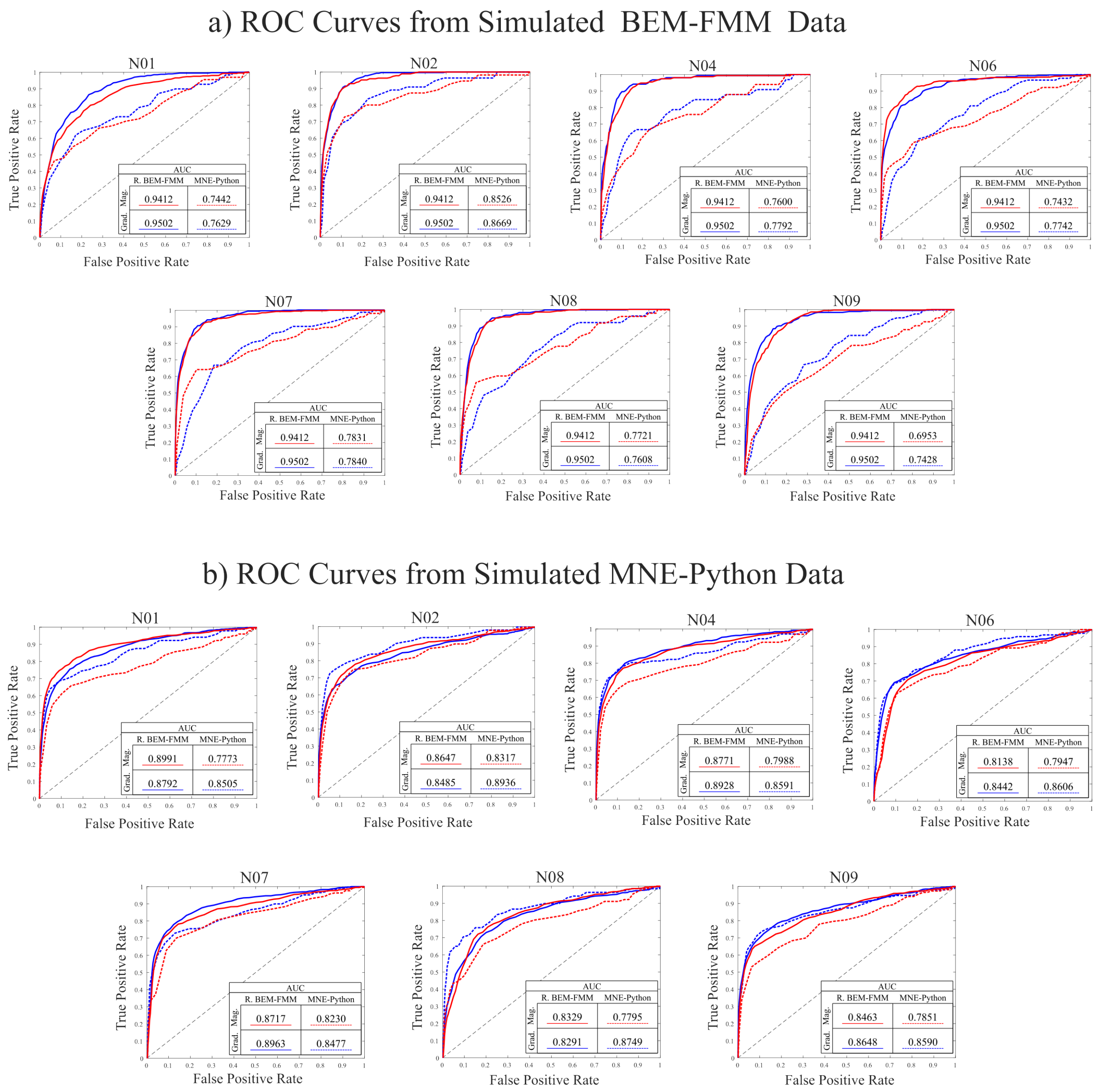


Figure 6: The ROC curves across all participants for the reciprocal BEM-FMM and MNE-Python models form simulated magnetometer and gradiometer data created from a) direct BEM-FMM, corresponding to Figure 2 and Figure 3, and b) MNE-Python, corresponding to Figure 4 and Figure 5.

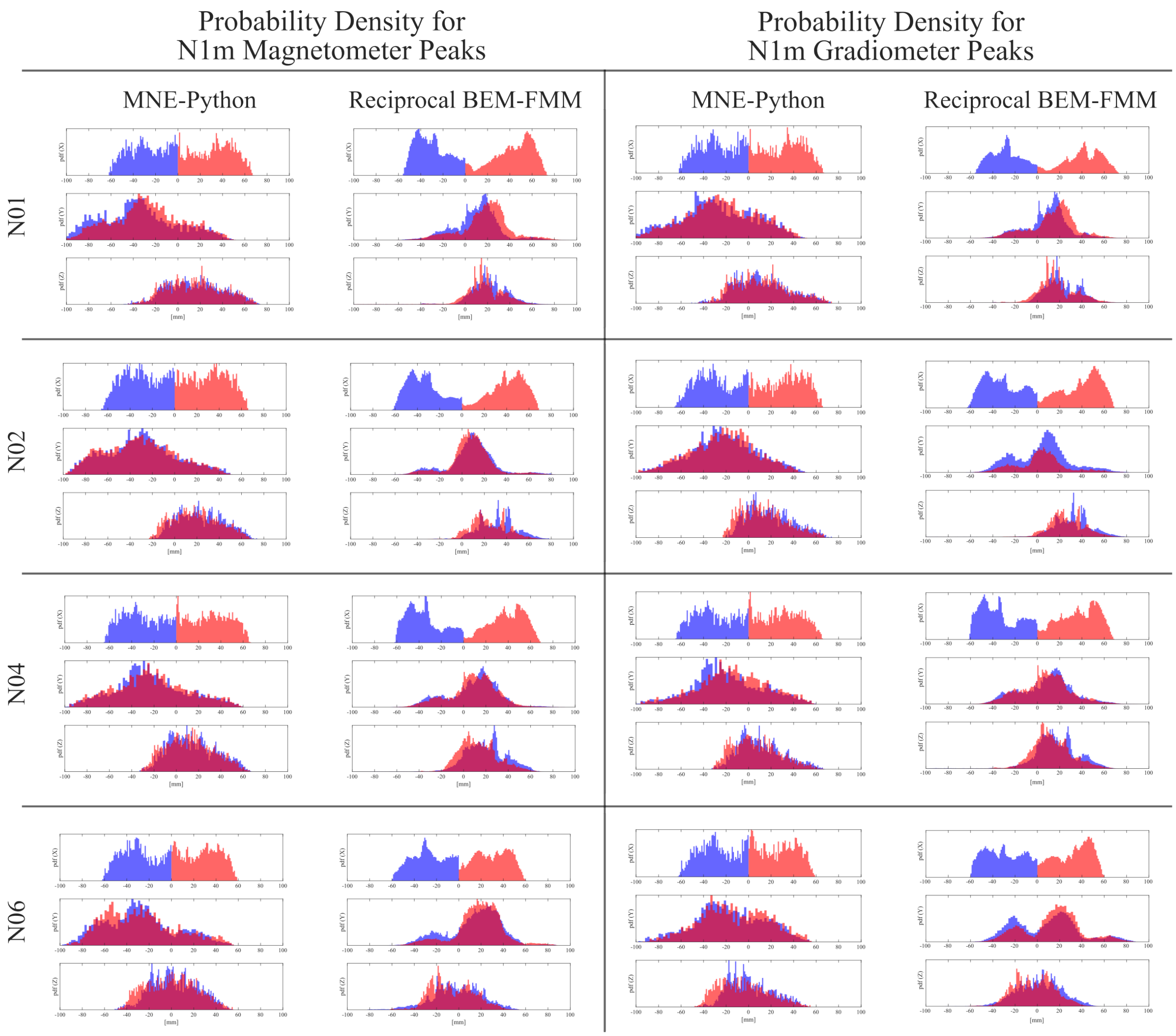


Figure 7: The probability distributions corresponding to each of the source localization solutions from participants N01, N02, N04, and N06 given in Figures 2 and 3 of the main paper. Each plot shows the distribution with respect the Cartesian axis directions. Blue indicates the left hemisphere, pink indicates the right hemisphere, and magenta is the intersection of the distributions.

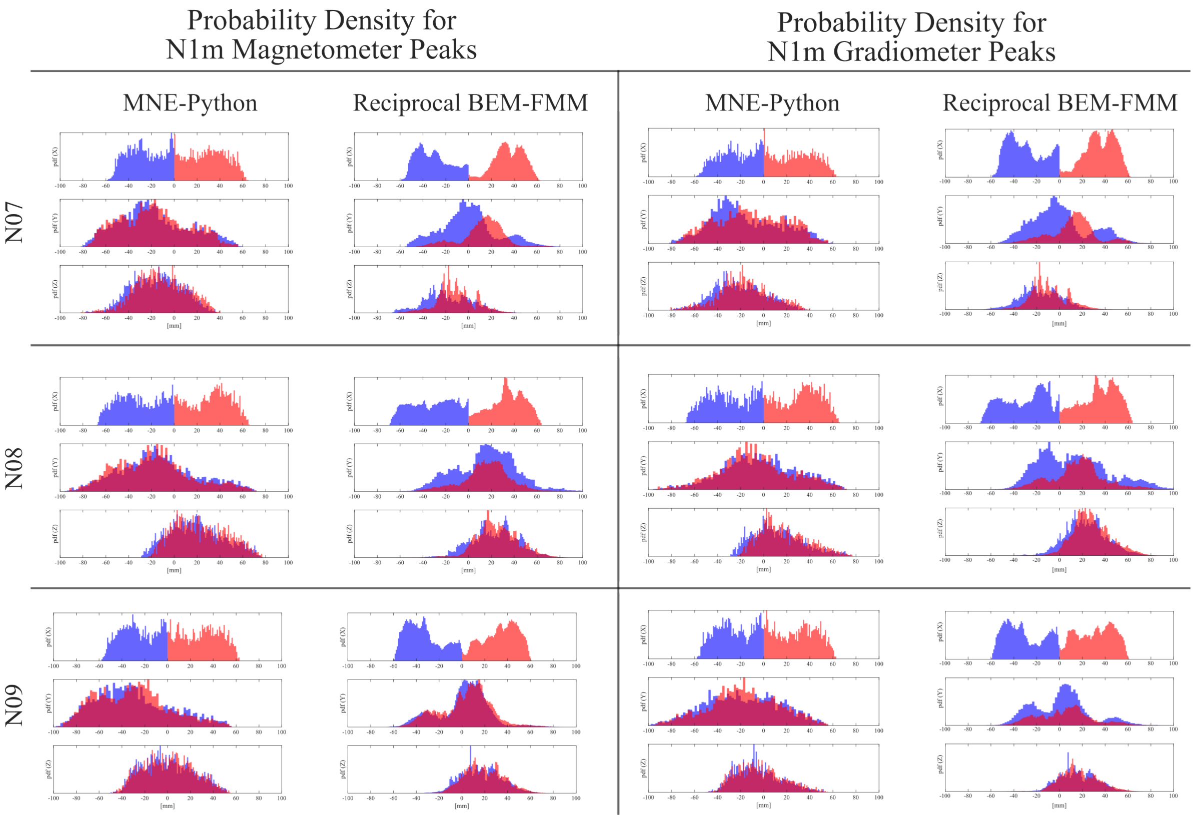


Figure 8: The probability distributions corresponding to each of the source localization solutions from participants N07, N08, and N09 given in Figures 2 and 3 of the main paper. Each plot shows the distribution with respect the Cartesian axis directions. Blue indicates the left hemisphere, pink indicates the right hemisphere, and magenta is the intersection of the distributions.
